# Using the simple telegraph model to decipher transcriptional burst regulation across genome-wide data

**DOI:** 10.1101/2025.10.29.685305

**Authors:** Liang Chen, Yuwen Wu, Chengkai Yang, Sijia Fang, Yu Liao, Yueheng Wu, Guozhi Jiang, Jianshe Yu, Feng Jiao

**Author notes:** Correspondence: G. Jiang, J. Yu, F. Jiao.

## Abstract

Gene transcription is a stochastic bursting process, with burst frequency and burst size as core parameters. Deciphering its genome-wide regulation is challenging, as complex models (capturing detailed biology) are impractical for large-scale use due to unstable inference and high computation. We proposed a simple telegraph model-based framework to analyze genome-wide scRNA-seq and smFISH datasets, addressing these limitations. When sample size ≥ 500 and burst parameter change ≥ 3-fold, its inferred burst frequency- and size-dominated variations reliably proxy true regulation. Analyses across mouse cells (CAST/C57 alleles, fibroblasts vs. embryonic stem cells) and healthy/hypertrophic cardiomyopathy (HCM) human heart tissues revealed key rules: 1) Burst frequency- or size-dominated regulation was primary (over 70% of variable genes), with more bursty-regulated genes in HCM; 2) TATA-initiator synergy (enhancing burst size-dominated regulation) was lost in HCM; 3) Burst frequency-dominated genes enriched in genome stability/cell cycle/apoptosis (via TFs like *Foxo4*/*Mcm2*), while burst size-dominated ones in signaling/metabolism/proteostasis (via TFs like *Zfp322a*/*Smchd1*/*Ppargc1a*), and their interdependent dysregulation accelerated HCM. This study establishes the simple telegraph model as a scalable framework linking transcriptional burst dynamics to cell fate and pathology.

## Introduction

Gene transcription is an inherently stochastic process, with messenger RNA (mRNA) synthesis across nearly all active genomic loci characterized by irregular inactive periods [1–3]. These observations confirm that gene transcription is not a continuous process but instead exhibits the hallmark of random bursting. This bursting pattern involves random initiation at discrete time points, the generation of substantial mRNA molecules during active periods, and subsequent transitions back to an inactive phase. Such transcriptional bursting directly drives fluctuations in mRNA copy numbers at the single-cell level, even within genetically homogeneous (isogenic) cell populations [4, 5]. To quantitatively characterize the dynamics of transcriptional bursting, two core parameters are universally employed: burst frequency, which measures the average rate at which transcription switches from the inactive to the active phase, and burst size, representing the average number of mRNA molecules generated per active period of a gene [2, 6]. Accurate estimation of burst frequency and size, along with elucidation of how these parameters are regulated in response to changes in the cellular environment, is of significant importance for various research fields. These include genetic engineering in cell stress responses [5, 7], exploration of the molecular mechanisms underlying cell fate determination [3, 8], and identification of therapeutic targets for diseases [9, 10].

In single cells, RNA labeling and visualization techniques, when coupled with input signal control, enable real-time imaging of transcriptional dynamics with high sensitivity and resolution, capturing transient events such as transcriptional burst frequency and size [1, 11]. Advanced versions of these methods further integrate next-generation sequencing, thereby achieving genome-wide detection at single-nucleotide resolution [2]. However, these imaging-based strategies are inherently time-consuming and require inserting repetitive sequences into target genetic loci, and exhibit strict dependence on viable, metabolically active cells. These limitations collectively restrict their widespread application to endogenous genes in mammalian cells across diverse sample types. To overcome these challenges, large-scale steady-state datasets on mRNA distributions in individual cells have been generated using single-cell RNA sequencing (scRNA-seq) and single-molecule fluorescence in situ hybridization (smFISH) [6, 8, 12, 13]. While these approaches cannot directly visualize real-time transcriptional dynamics, they circumvent this limitation by employing mathematical models to infer key transcriptional parameters, including burst frequency and burst size, from genome-wide mRNA expression profiles across sampled single cells.

Stochastic gene transcription is typically modeled using the telegraph model, which employs first-order rate constants: *k*_on_ for gene activation, *k*_off_ for inactivation, and *k*_b_ for mRNA synthesis in the active state [4, 6, 8] (Fig. 1a). Recent advancements in single-cell techniques now allow precise measurements of mRNA copy numbers across hundreds to thousands of cells, yielding extensive transcriptional distribution datasets. The telegraph model is widely applied to fit these data, enabling the estimation of transcriptional burst frequency *b*_*f*_ = *k*_on_ and burst size *b*_*s*_ = *k*_b_*/k*_off_ under various induction conditions or promoter architectures. Notably, transcriptional upregulation follows diverse strategies: increasing burst frequency of *Oct4* and *Nanog* during the cell-cycle of mouse embryonic stem cells [14]; elevating burst size by reducing *k*_off_ for over 20 *E. coli* promoters [15], or by increasing *k*_b_ for the serum-induced mammalian *ctgf* gene [16]. Co-regulation of both parameters also occurs [5, 7]. For instance, the HIV LTR promoter in human T lymphocytes shows distinct regulatory patterns across different integration sites: burst frequency dominates when gene expression is low, whereas burst size becomes the primary regulatory target at high expression levels [7].

**Figure 1.**
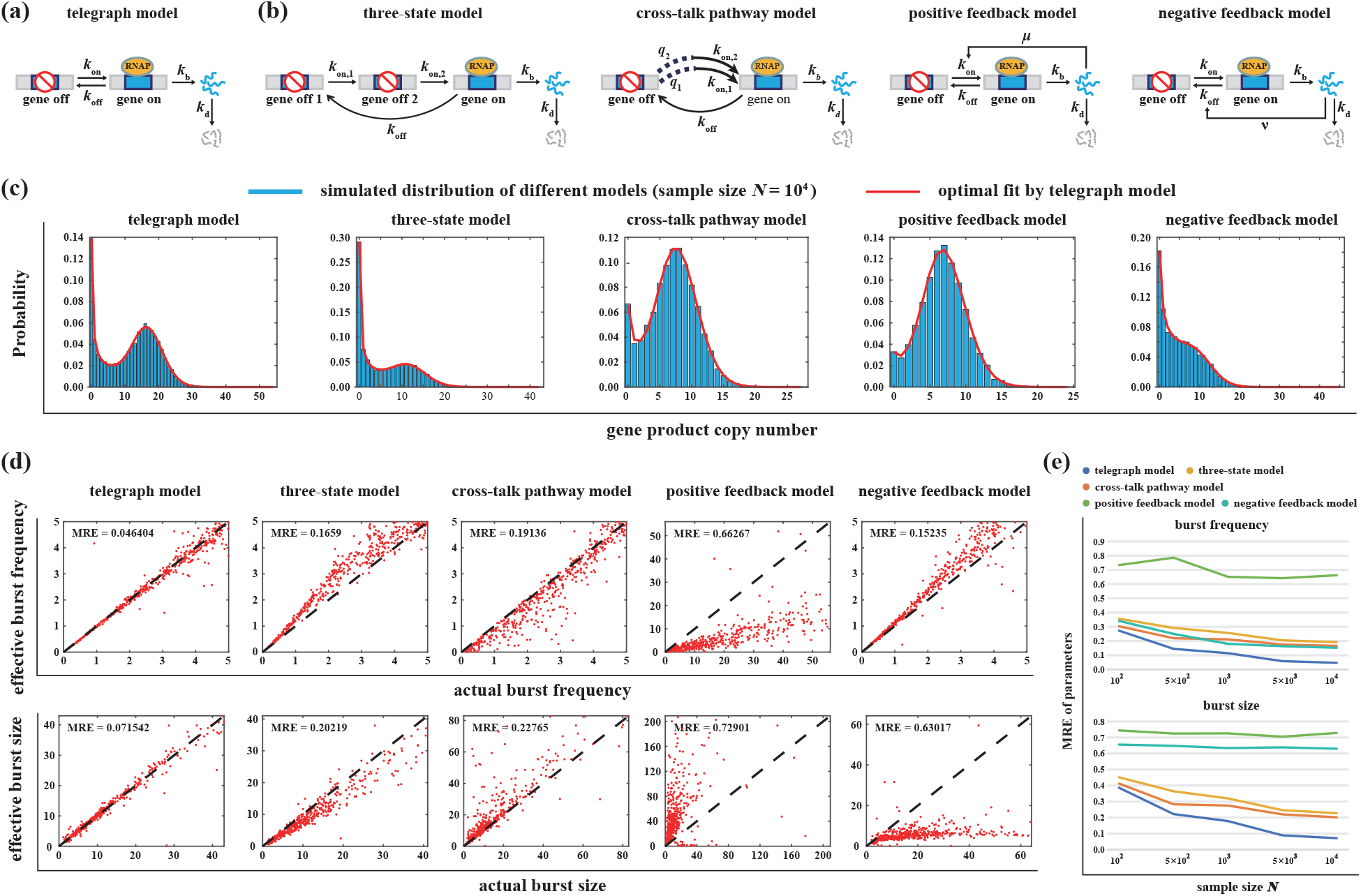
Evaluation of the simple telegraph model in fitting complex gene transcription models. **(a)** Diagram illustrating the telegraph model with activation rate *k*_on_, inactivation rate *k*_off_, and synthesis rate *k*_b_; The model defines effective burst frequency *b*_*f*_ = *k*_on_ and burst size *b*_*s*_ = *k*_b_*/k*_off_. **(b)** Diagram illustrating the three-state model (with refractory inactive periods), cross-talk pathway model (competitive TF binding), positive and negative feedback models (autoregulatory circuits). **(c)** Steady-state gene product copy number distributions (SSA-simulated) for each model; all are well approximated by the effective telegraph model (MLE-fitted). **(d)** Scatter plots comparing estimated effective *b*_*f*_ and *b*_*s*_ with their true burst frequency and burst size across 500 parameter sets, under large sample size *N* = 10^4^. The telegraph model accurately captures its own parameters (mean relative error, MRE < 0.2), while most complex models exhibit significant biases (MRE > 0.2). **(e)** MRE of *b*_*f*_ and *b*_*s*_ estimates across sample sizes (*N* = 10^2^, 5 *×* 10^2^, 10^3^, 5 *×* 10^3^, 10^4^). MRE decreases with increasing *N* for all models; *b*_*f*_ estimation is more accurate than *b*_*s*_ estimation; no complex model achieves MRE < 0.2 for both *b*_*f*_ and *b*_*s*_ (even at *N* = 10^4^), and the telegraph model shows MRE > 0.2 at small *N* = 10^2^.

The traditional telegraph model has significant limitations, as it neglects key biological mechanisms that influence transcriptional dynamics. These mechanisms include the multistep recruitment of transcription factors (TFs), pathway competition during gene activation [17, 18], and feedback regulation, such as the involvement of over 40% of *E*.,*coli* TFs in negative feedback networks [19] and the HIV Tat-mediated positive feedback loop associated with HIV latency [20]. Incorporating these detailed mechanisms into the telegraph model has yielded more complex extended frameworks (Fig. 1b). A critical challenge arises in model selection and parameter inference for these extended models. Unlike the well-established analytical framework for the telegraph model, conventional methods for analyzing complex models are both computationally intensive and unreliable when applied to typical scRNA-seq or smFISH datasets [21, 22]. Furthermore, several proposed strategies to differentiate between specific models often rely on temporal data and are tailored to particular model types, thereby limiting their ability to compare across a broad range of complex frameworks [23–25]. We recently introduced an alternative approach: fitting steady-state datasets generated by complex models using the simple telegraph model [21]. However, this method requires elaborately designed genetic circuits and measurements under multiple induction conditions, which limits its generalization to genome-wide datasets.

To overcome limitations of the classical telegraph model (neglecting complex regulation) and impracticalities of complex models for genome-wide analysis (unstable inference, large-sample dependence), we employed the simple telegraph model across genome-wide scRNA-seq and smFISH datasets. Though the model exhibited quantitative biases when estimating burst frequency *b*_*f*_ and size *b*_*s*_ for genes under complex regulation, it reliably captures qualitative burst dynamics. Specifically, with a cell sample size *N* ≥ 500 and a 3-fold change in the model’s *b*_*f*_ or *b*_*s*_, the inferred *b*_*f*_ - or *b*_*s*_-dominated variations serve as robust proxies for true regulatory changes. We further identified three context-conserved rules governing burst regulation, validated across mouse fibroblasts versus embryonic stem (ES) cells (CAST/C57 alleles) and healthy versus hypertrophic cardiomyopathy (HCM) human heart tissues. These rules encompass: 1) primary transcriptional burst regulatory modes, 2) synergistic effect of promoter TATA and initiator elements and its loss in HCM, and 3) functional specialization of these regulatory modes-such modes form a vicious cycle in HCM to accelerate myocardial degeneration. These findings lay a foundation for understanding transcriptional burst control and its pathological roles.

## Results

### Quantifying bias in parameter inference: Using the telegraph model to fit complex models

In genome-wide transcriptional analysis, assigning a unique model for each gene’s transcriptional dynamics is impractical. A more efficient alternative is to fit genome-wide datasets using the simple telegraph model (Fig. 1a), thereby inferring two core transcriptional parameters: burst frequency *b*_*f*_ and burst size *b*_*s*_ [6, 13]. However, this approach raises two critical questions. First, for genes whose transcription is governed by complex regulatory frameworks rather than the telegraph model’s simple two-state dynamics, how do inferred *b*_*f*_ and *b*_*s*_ deviate from the true kinetic rates of the gene’s native regulation? Second, how does sample size-number of cells in genome-wide datasets-influence the discrepancy between inferred parameters and real kinetic rates? Notably, sample size can introduce non-negligible noise into steady-state gene product distribution data, thereby skewing parameter estimation [22]. Consequently, even for genes strictly following the telegraph model’s regulatory principles, model-fitted *b*_*f*_ and *b*_*s*_ may deviate from their true values.

Here, we examine four commonly used complex models (Fig.,1b). The three-state model posits that inactive period gene involve two rate-limiting steps and “refractory” behavior, reflecting multi-step promoter activation [26]. This model has been successfully applied to explain the non-exponential, peaked inactive period distributions observed in some genes in mammalian and bacterial cells [11, 27]. Notably, the telegraph and three-state models describe only a single gene activation pathway. In contrast, the cross-talk pathway model captures scenarios where real gene activation involves multiple signaling pathways [28]. This model characterizes competitive TFs binding to the promoter, and has been used to reproduce rapid mRNA overshoot in tumor necrosis factor-induced mouse fibroblasts [29, 30] and arabinose/isopropylthio-L-D-galactoside-induced *E. coli* [21]. The positive/negative feedback model describes an autoregulatory circuit, where the gene’s encoded protein either activates (positive feedback) or represses (negative feedback) its own expression. Its reaction scheme mirrors the telegraph model, with the key distinction being that the protein modulates gene activity via a feedback strength rate constant [31, 32]. Exact formulas for the gene product distributions of these models are provided in the Supplementary Materials, and calculations of burst frequency and size are detailed in the Methods and Materials section.

For each of the five stochastic gene transcription models under investigation, we first generated 500 independent parameter sets (Methods and Materials). For each combination of model and parameter set, we employed the Stochastic Simulation Algorithm (SSA) to simulate synthetic gene product data across five cell population sizes: *N* = 10^2^, 5 × 10^2^, 10^3^, 5 × 10^3^, 10^4^ cells. Subsequently, we applied maximum-likelihood estimation (MLE) [6, 21] to fit the simulated steady-state distributions data of gene products to the telegraph model (Methods and Materials). As shown in Fig. 1c: consistent with the telegraph model, each of the four more complex models generated either unimodal or bimodal gene product distributions [33, 34]. Importantly, the steady-state distributions simulated by the four complex models could generally be well approximated by the telegraph model [21]. From this fitting, we inferred two effective parameters of the telegraph model: the effective burst frequency *b*_*f*_ = *k*_on_ and the effective burst size *b*_*s*_ = *k*_b_*/k*_off_.

For each model and sample size tested, we systematically analyzed 500 independent parameter sets. We generated scatter plots where each data point represented a pair of values: the estimated effective *b*_*f*_ (or *b*_*s*_) versus their corresponding true values from the parameter sets. To assess estimation accuracy, we calculated the deviation of each data point from the 1:1 reference line, which indicates perfect agreement between estimated and true values. We quantified this deviation using the mean relative error (MRE) between the 500 estimated value pairs and the 1:1 reference line, with representative results shown in Fig. 1d.

At large sample size of *N* = 10^4^, the telegraph model demonstrated high accuracy, with its *b*_*f*_ (or *b*_*s*_) estimates clustered tightly around the 1:1 reference line (MRE < 0.2), confirming its reliability in capturing true *b*_*f*_ and *b*_*s*_ dynamics. In contrast, for other complex models, the 500 data point for *b*_*f*_ and *b*_*s*_ generally fell either above or below the 1:1 line. Points above the line indicated overestimation of the effective *b*_*f*_ or *b*_*s*_, while points below suggested underestimation. Notably, the three-state, cross-talk pathway and negative feedback models showed close alignment between effective *b*_*f*_ and their true values (MRE < 0.2). However, for all complex models, effective *b*_*s*_ estimates exhibited significant deviations from the 1:1 reference line, with MRE values exceeding 0.2. This indicates a notable bias in parameter estimation for these models relative to their true values.

To investigate the impact of sample size *N* on the accuracy of estimated effective *b*_*f*_ and *b*_*s*_, we analyzed MRE values across *N* = 10^2^, 5 × 10^2^, 10^3^, 5 × 10^3^, 10^4^ for each model (Fig. 1e, Supplementary Fig. S1). This analysis revealed three key patterns. First, across all models, MRE for both *b*_*f*_ and *b*_*s*_ consistently decreased as *N* increased, aligning with statistical principles where larger sample sizes reduce estimation uncertainty. Additionally, *b*_*f*_ MRE was consistently smaller than *b*_*s*_ MRE for every model at each sample size, confirming that *b*_*f*_ estimation is more accurate than *b*_*s*_, consistent with previous findings [21, 36]. Second, even at the largest *N* = 10^4^, no complex models achieved MRE < 0.2 for both *b*_*f*_ and *b*_*s*_ simultaneously. This underscores that the simple telegraph model cannot quantitatively capture the regulatory dynamics of real biological systems. Third, under the smallest *N* = 10^2^, MRE for both *b*_*f*_ and *b*_*s*_ exceeded 0.2 even for the telegraph model. This highlights that the telegraph model fails to accurately reflect its inherent regulatory properties under small samples, likely due to non-negligible noise from limited data overwhelming its ability to distinguish true parameter signals from random variation.

### The telegraph model reliably reflects burst frequency- and burst size-dominated regulation

Previous findings have shown that the telegraph model exhibits significant quantitative biases in estimating burst frequency and size. Here, we further investigate whether this simple telegraph model can still qualitatively capture changes in burst frequency and size across different model configurations. To address this question, we designed the following analysis: For each model and each parameter under investigation, we generated 500 random pairs of parameter sets. Within each pair, the target parameter was set to two distinct values while all other parameters remained identical. This design simulates scenarios where a single parameter is altered under two conditions (different cellular environments [7] or promoter configurations [5]), with all other variables held constant.

Here, we outline the analytical workflow: First, we generated synthetic distribution data using the original model with two distinct parameter values for each parameter pair. Next, we applied the telegraph model to fit these synthetic datasets, obtaining estimated values for the effective burst frequency *b*_*f*_ and size *b*_*s*_ under each condition. To systematically evaluate the qualitative consistency between estimates and true values, we computed two types of ratios for each pair: 1) the ratio of estimated *b*_*f*_ (or *b*_*s*_) values between the two conditions, and 2) the ratio of their corresponding ground-truth values derived from the original model. These ratio pairs were then plotted against each other (Fig. 2). To enhance clarity, we grouped the data into bins of 20 genes each, calculated the rolling median for ratios within each bin [6], and quantified the strength of association between estimated and ground-truth ratios using the Kendall’s tau correlation coefficient (KCC). This comprehensive approach allows us to rigorously assess whether the telegraph model captures both the direction (e.g., increase vs. decrease) and relative magnitude of changes in burst characteristics, despite its known quantitative biases.

**Figure 2.**
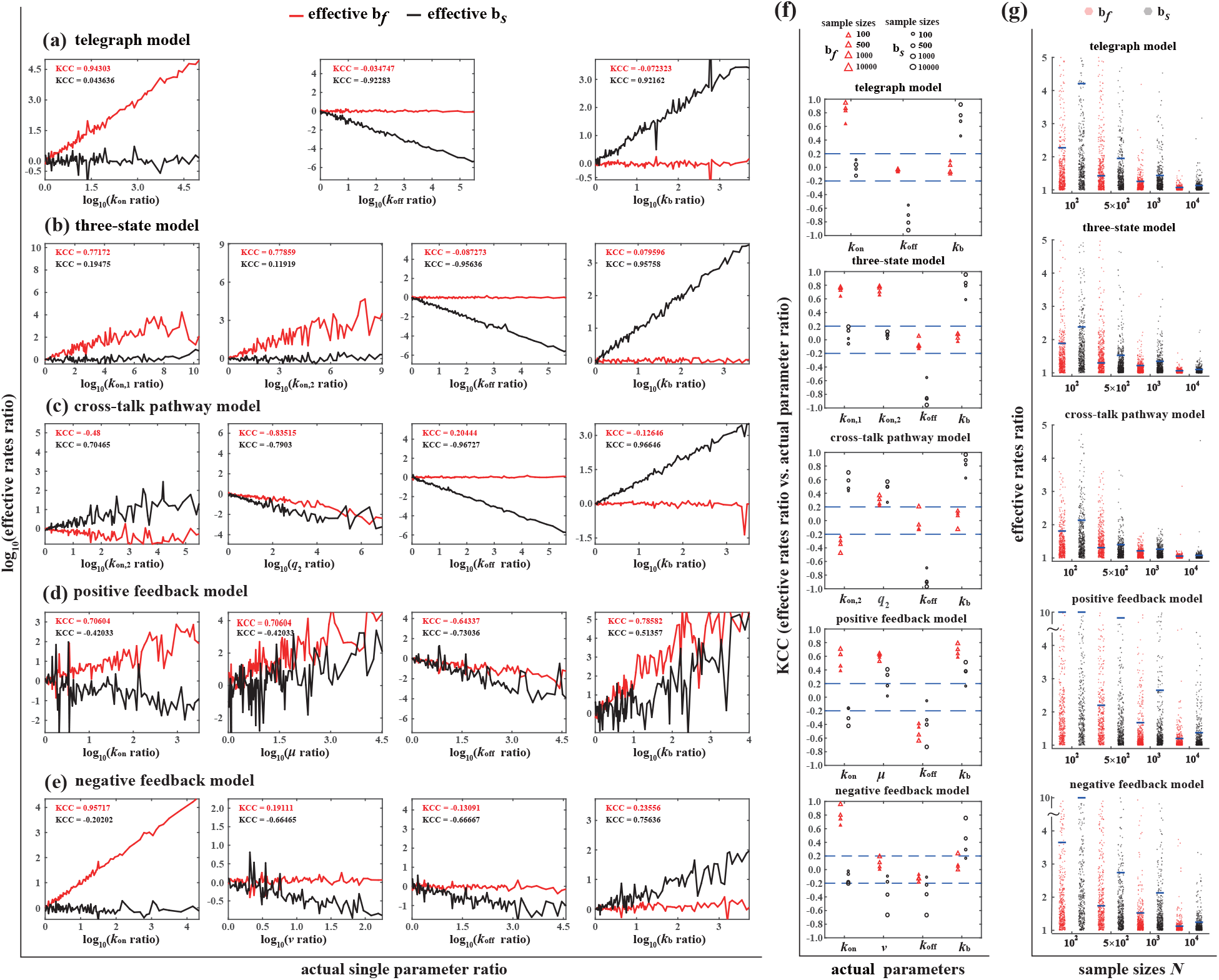
Qualitative reliability of the telegraph model in capturing burst frequency and size variations and threshold determination. **(a)-(e)** Under cell sample size *N* = 10^4^: For the telegraph model (a) and four complex models ((b)-(e): three-state, cross-talk pathway, positive feedback, negative feedback), 500 independent parameter pairs were generated per model by varying one target parameter while keeping others fixed. Synthetic steady-state gene product distribution data were generated and fitted by the telegraph model to estimate effective *b*_*f*_ (burst frequency) and *b*_*s*_ (burst size). Scatter plots of ratios of effective vs. true *b*_*f*_ (or *b*_*s*_) were constructed, with Kendall’s tau correlation coefficients (KCC) indicating tracking ability. Results: effective *b*_*f*_ - or *b*_*s*_-dominated variations correspond to true burst frequency- or size-dominated regulation. **(f)** Stability of KCC for *b*_*f*_ (or *b*_*s*_) ratios across sample sizes *N* for each model and its varying parameters: KCC values of *b*_*f*_ (or *b*_*s*_) ratios at *N* = 10^4^ (largest) were categorized into three intervals: (*−*0.2, 0.2), (*−*1, 0.2), (0.2, 1). A variation pattern was stable if KCC remained in the same interval across smaller *N*, and disrupted if shifted. Results: telegraph, three-state, and cross-talk pathway models show stable patterns across all *N* ; positive/negative feedback models are disrupted at *N* = 10^2^, but regain stability at *N ≥* 500. **(g)** Fold-change thresholds for *b*_*f*_ and *b*_*s*_: For each *N* and model, 500 random parameter sets were generated; two synthetic data groups per set were fitted by the telegraph model to obtain *b*_*f*_ and *b*_*s*_. The ratio of the larger to smaller *b*_*f*_ (or *b*_*s*_) was computed, and the 75th percentile of these 500 ratios was defined as the threshold. For *N ≥* 500, a fold change in *b*_*f*_ or *b*_*s*_ exceeding a threshold of 3 indicates genuine regulatory changes, rather than sampling noise.

We first examine the largest sample size (*N* = 10^4^). For synthetic data from the telegraph model (Fig. 2a), the effective burst frequency *b*_*f*_ and size *b*_*s*_ reliably track variations in their true values, demonstrating the basic validity of these metrics. This pattern holds in the three-state model (Fig. 2b): variations in activation rates (*k*_on,1_, *k*_on,2_) induce *b*_*f*_ -dominated variation (marked changes in *b*_*f*_ with negligible *b*_*s*_ fluctuation), directly reflecting true frequency changes. Conversely, alterations in inactivation or synthesis rates (*k*_off_ or *k*_b_) produce *b*_*s*_-dominated variation (pronounced *b*_*s*_ shifts with stable *b*_*f*_), faithfully indicating real size variations. The cross-talk pathway model (Fig. 2c) exhibits mixed behavior. Modifications to pathway selection probabilities (*q*_1_, *q*_2_) or activation strengths (*k*_on,1_, *k*_on,2_) trigger co-variation (simultaneous changes in both *b*_*f*_ and *b*_*s*_), which do not map uniquely to true burst frequency variation. However, perturbations to inactivation or synthesis rates (*k*_off_ or *k*_b_) maintain *b*_*s*_-dominated patterns, reliably signaling true size changes. Feedback models exhibit distinct regulatory characteristics: positive feedback (Fig. 2d) results in coordinated co-variation of *b*_*f*_ and *b*_*s*_ with any parameter perturbation, offering no specific insight into underlying frequency or size variations. Negative feedback (Fig.,2e), however, preserves a clear separation: changes in activation rate *k*_on_ drive *b*_*f*_ -dominated variation (indicating true frequency indicators), while perturbations to feedback/inactivation/synthesis parameters (*ν/k*_off_*/k*_b_) induce *b*_*s*_-dominated effects (reliably indicating size changes).

Collectively, these findings across all models demonstrate that, effective *b*_*f*_ -dominated and *b*_*s*_-dominated variations of the telegraph model consistently serve as robust proxies for true frequency and size changes, respectively. In contrast, concurrent co-variation of *b*_*f*_ and *b*_*s*_ (observed in cross-talk pathway models under certain perturbations and universally in positive feedback systems) does not allow reliable inference about specific real-system variations.

Here, we evaluated the robustness of the variation patterns of the effective parameters *b*_*f*_ and *b*_*s*_ across different cell sample sizes *N* = 10^2^, 5 × 10^2^, 10^3^, 5 × 10^3^, 10^4^. For each model and each of its varying parameters, we first analyzed the largest sample size (*N* = 10^4^; see Fig. 2a-e) and categorized the KCC values of the corresponding *b*_*f*_ (or *b*_*s*_) ratio into three predefined intervals: (−0.2, 0.2), (−1, −0.2) or (0.2, 1). To assess stability, we compared these KCC values with those obtained under smaller sample sizes. A variation pattern of *b*_*f*_ (or *b*_*s*_) was deemed stable across the sample size range if its KCC value remained within the same interval as observed for *N* = 10^4^. Conversely, a shift to a different interval under smaller *N* indicated a disrupted pattern for those sample sizes.

Fig. 2f presents KCC values of *b*_*f*_ (or *b*_*s*_) ratios for each model and varying parameter. Our results reveal that *b*_*f*_ and *b*_*s*_ variation patterns are stable across all tested sample sizes for synthetic data from the telegraph, three-state, and cross-talk pathway models. For the positive and negative feedback models, nearly all varying parameters exhibited disrupted patterns under the smallest sample size *N* = 10^2^, except for the varying activation rate *k*_on_ in the negative feedback model. Notably, stability was restored for both feedback models when *N* ≥ 5 × 10^2^. These findings confirm that cell samples with *N* ≥ 5 × 10^2^ ensure robust prediction of real burst dynamics using the variation patterns of effective *b*_*f*_ and *b*_*s*_.

In studying steady-state gene product distributions, even when using identical models and parameters, limited cell samples can introduce non-negligible noise. This noise may lead to substantial differences in estimated effective parameters like *b*_*f*_ or *b*_*s*_ between datasets-discrepancies that may arise from sampling noise rather than genuine biological regulatory changes. This introduces a critical interpretive challenge: in real experiments, inferred *b*_*f*_ - and *b*_*s*_-dominated regulation may reflect sampling noise from small samples rather than true transcriptional differences between cellular environments. To address this, defining a fold-change threshold for *b*_*f*_ (or *b*_*s*_) is essential. When the observed fold change exceeds this threshold, the inferred regulatory dominance is more likely due to genuine transcriptional changes; below it, the difference is more likely attributable to sampling noise. This threshold helps distinguish true biological signals from sampling artifacts.

To define the threshold, we conducted a theoretical analysis as follows: For each sample size and each model, we generated 500 random parameter sets. For each set, we produced two groups of synthetic data, which were then fitted using the telegraph model to yield two corresponding values for *b*_*f*_ (or *b*_*s*_) per group. We subsequently calculated the ratio of the larger *b*_*f*_ (or *b*_*s*_) to the smaller one, resulting in 500 such ratios (all ≥ 1) for *b*_*f*_ (or *b*_*s*_) per sample size and model. The cutoff for *b*_*f*_ (or *b*_*s*_) is defined as the 75th percentile of these 500 ratios. As shown in Fig. 2g, the thresholds for both *b*_*f*_ and *b*_*s*_ decrease as sample size increases, with the *b*_*f*_ threshold consistently lower than that of *b*_*s*_. This pattern highlights two important insights: first, larger sample sizes enhance the reliability of identifying whether regulation is dominated by effective burst frequency or burst size; second, conclusions about frequency-dominated regulation tend to be more reliable than those regarding burst size-dominated regulation.

To apply these thresholds to genome-wide data analysis using the telegraph model, we focused on scenarios involving effective *b*_*f*_ -dominated variation and *b*_*s*_-dominated variation, as these two types of variation reflect genuine changes in the system’s bursting properties (Fig. 2a-e). Consequently, we excluded the positive feedback model, as variations in its parameters lead to co-variation of effective *b*_*f*_ and *b*_*s*_, complicating interpretation. We restricted our analysis to sample sizes exceeding *N* = 10^2^, as Fig. 2f indicates that effective *b*_*f*_ -dominated and *b*_*s*_-dominated variations at a sample size of 100 may not reliably capture true bursting changes. Within the range of 500 to 10^4^ samples, the negative feedback model exhibited the highest thresholds for for *b*_*f*_ and *b*_*s*_ among the telegraph, three-state, and cross-talk pathway models. Specifically, for 500 and 1000, the *b*_*f*_ threshold remained below 2, and the *b*_*s*_ threshold below 3; at *N* = 10^4^, both thresholds were within 2. These threshold determinations carry the following implication: when using effective *b*_*f*_ -dominated or *b*_*s*_-dominated variation to infer genuine system-level regulation dominated by burst frequency or size, a fold change in effective *b*_*f*_ or *b*_*s*_ exceeding 3 provides robust evidence of significant changes in the corresponding parameters of the real system. Such changes are unlikely to be solely attributable to noise related to sample size.

### Transcriptomic regulation of burst frequency and size in mouse fibroblasts versus ES cells

To investigate transcriptome-wide patterns of transcriptional bursting, we analyzed the kinetic parameters *b*_*f*_ and *b*_*s*_, previously inferred using the MLE method for 7,186 genes [6]. These parameters were derived from transcriptomic data of individual primary mouse fibroblasts and mouse ES cells, with each allele (CAST/EiJ × C57BL/6J) analyzed separately. Moving forward, we will focus on 4,854 genes of C57 allele and 4,769 genes of CAST allele shared between fibroblasts and ES cells (i.e., expressed with inferable kinetics) to determine the differences in their bursting parameters *b*_*f*_ and *b*_*s*_ (Table S1). Building on our previous findings, we categorized the variations in *b*_*f*_ and *b*_*s*_ into four patterns: *b*_*f*_ -dominated variation (where *b*_*f*_ changes by more than 3-fold while *b*_*s*_ changes by less than 3-fold), *b*_*s*_-dominated variation (where *b*_*s*_ changes by more than 3-fold while *b*_*f*_ changes by less than 3-fold), co-variation of *b*_*f*_ and *b*_*s*_ (both parameters changing by more than 3-fold), and no significant transcriptional changes (both parameters changing by less than 3-fold).

For transcriptional changes in the C57 allele between mouse fibroblasts and ES cells, 61.2% of genes exhibited no significant changes in either *b*_*f*_ or *b*_*s*_, indicating minimal transcriptional differences. Among the remaining 40% of genes with significant changes in either *b*_*f*_ or *b*_*s*_, *b*_*f*_ -dominated variation was the most prevalent (17.4%), followed by *b*_*s*_-dominated variation (11.6%). Our theory suggests that these gene-specific variations most likely reflect genuine burst frequency- or size-dominated regulatory mechanisms between the two cell types (see Fig. 3a). To further explore the relationship between bursting parameters and gene expression levels, we plotted the ratio of *b*_*f*_ (or *b*_*s*_) between the two cell types against the ratio of their average expression levels for each gene. Both *b*_*f*_ ratios and *b*_*s*_ ratios exhibited positive correlations with the average expression ratio, with the correlation between *b*_*f*_ and average expression being more pronounced, consistent with previous findings [6]. Notably, transcriptional bursting in the CAST allele displayed nearly identical regulatory characteristics to those observed in the C57 allele (see Fig. 3a).

**Figure 3.**
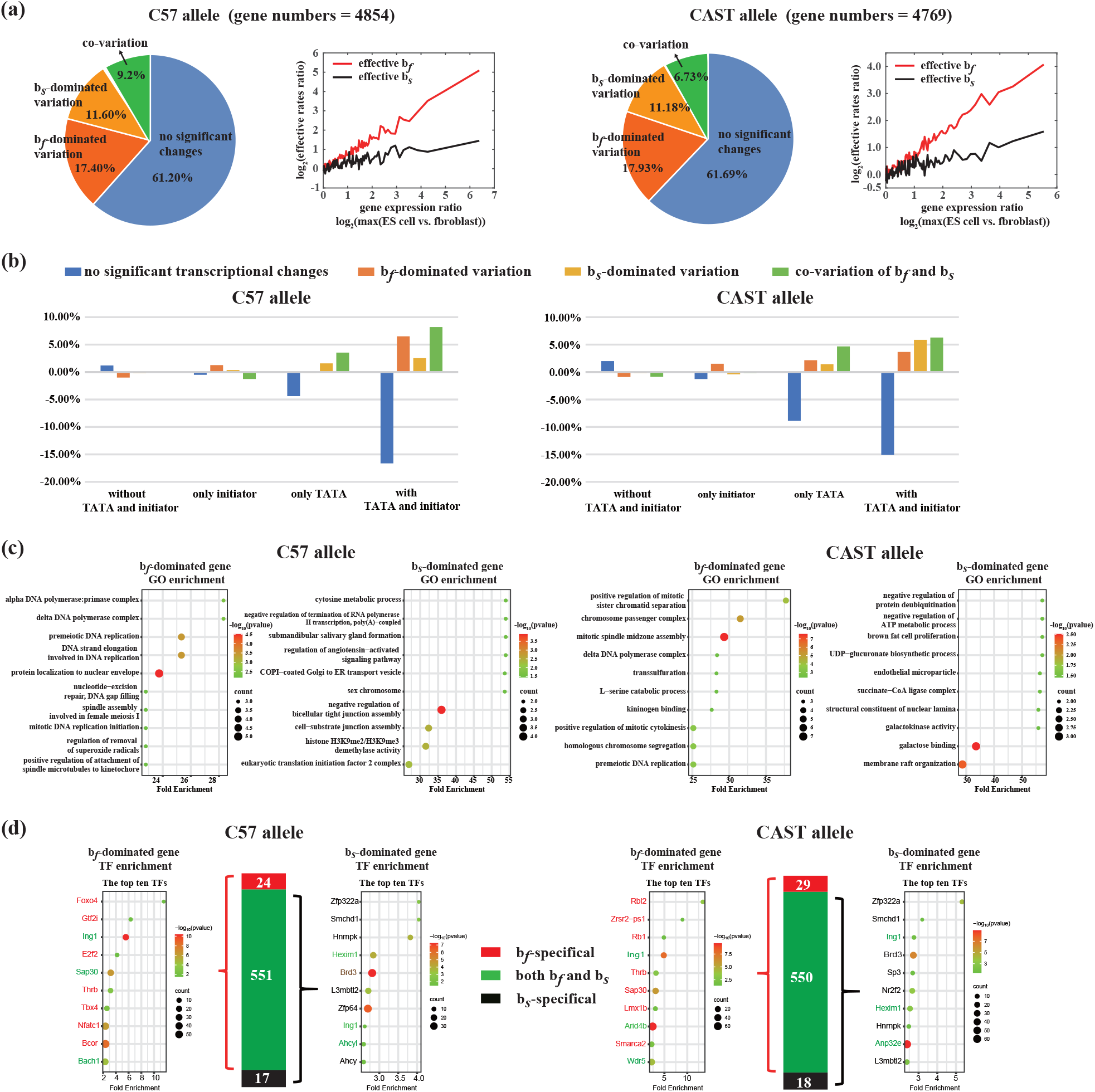
Transcriptional burst regulation in mouse fibroblasts vs. ES cells (CAST/C57 alleles) **(a)** Effective *b*_*f*_ (burst frequency) and *b*_*s*_ (burst size) variation for genes shared between mouse fibroblasts and ES cells (C57 allele: 4854 genes; CAST allele: 4769 genes [6]). Genes were categorized into four types: *b*_*f*_ -dominated, *b*_*s*_-dominated, co-variation, and no significant change. For both C57 and CAST, *b*_*f*_ -dominated (17.4%) and *b*_*s*_-dominated (11.6%) variations were the primary types of significant change. Insets show positive correlations between *b*_*f*_ (or *b*_*s*_) ratios and average gene expression ratios, with a stronger correlation for *b*_*f*_. **(b)** Impact of promoter architectures (only TATA, only initiator, both, neither) on *b*_*f*_ and *b*_*s*_ variations: Proportions of each variation type in each promoter group were compared to the baseline (unstratified genes in (a)); positive/negative values indicate promotion/inhibition. Results: Initiator had no significant effect; TATA elements independently reduced the “no significant change” proportion; TATA-initiator synergy further enhanced *b*_*f*_ - and *b*_*s*_-dominated variations. **(c)** Functional enrichment of *b*_*f*_ - and *b*_*s*_-dominated variation genes (both alleles): *b*_*f*_ -dominated genes were enriched in GO terms related to cell division and genome stability; *b*_*s*_-dominated genes were enriched in signaling transduction, energy/material metabolism, cell structure/organelle organization. **(d)** Enriched TFs included specific (regulating only *b*_*f*_ or *b*_*s*_) and general (regulating both) types, with specific TFs accounted for 70% of top 10 TFs. Both alleles shared *b*_*f*_ -specific TF such as and *b*_*s*_-specific TFs such as *Zfp322a* and *Smchd1*.

We further investigated how TATA and initiator elements in promoters regulate *b*_*f*_ and *b*_*s*_. For both the C57 and CAST alleles, genes were first grouped by their promoter architectures: those with only TATA elements, only initiator elements, both elements, or neither; see Fig. 3b. Each promoter-based group was then categorized into the four types of *b*_*f*_ and *b*_*s*_ variations: no changes in *b*_*f*_ and *b*_*s*_; *b*_*f*_ -dominated variation; *b*_*s*_-dominated variation; and co-variation of both. To evaluate the regulatory impact of each promoter component, we calculated the proportion of genes in each *b*_*f*_ and *b*_*s*_ category within each promoter group (Table S2). This proportion was then subtracted from the corresponding baseline proportion of each *b*_*f*_ and *b*_*s*_ category (i.e., the proportion observed when genes were not stratified by promoter components in Fig. 3a). A positive value from this subtraction indicates that the specific promoter component promotes the corresponding *b*_*f*_ and *b*_*s*_ regulation, while a negative value indicates inhibition; minimal deviation implies little to no impact.

For both the C57 and CAST alleles (Fig. 3b), among genes where neither *b*_*s*_ nor *b*_*f*_ changed, promoters lacking both TATA and initiator elements, or containing only initiator elements, had no significant effect on promoting variations in *b*_*s*_ or *b*_*f*_. In contrast, promoters containing only TATA elements significantly reduced the proportion of genes in the “unchanged *b*_*f*_ and *b*_*s*_” category, demonstrating that the TATA element independently promotes variations in *b*_*f*_ and *b*_*s*_. Notably, promoters containing both TATA and initiator elements exhibited a marked synergistic effect: they drastically reduced the proportion of genes with unchanged *b*_*f*_ and *b*_*s*_, directly confirming that the combination of these two elements synergistically promotes variations in *b*_*f*_ and *b*_*s*_. Moreover, this synergism leads to distinct types of variation in different alleles: in the C57 allele, the synergistic effect primarily drives *b*_*f*_ -dominated variation, whereas in the CAST allele, it mainly promotes *b*_*s*_-dominated variation. In summary, the initiator element has no significant impact on variations in *b*_*f*_ or *b*_*s*_; the TATA element can independently promote such variations; and the synergistic interaction between these two promoter elements specifically drives either *b*_*f*_ -dominated or *b*_*s*_-dominated regulatory variations.

We focused on two groups of genes: the *b*_*f*_ -dominated variant genes and *b*_*s*_-dominated variant genes. Our results revealed that these two groups of genes are subject to frequency-dominated regulation and burst-dominated regulation, respectively. For the frequency-dominated regulated genes in both C57 and CAST alleles, nearly all enriched Gene Ontology (GO) terms clustered tightly around cell division processes, which are critical for maintaining genomic stability and division fidelity. These processes include the initiation and elongation of DNA replication, the repair of replication damage and oxidative damage, DNA replication prior to mitosis, and the regulation of spindle assembly and chromosome-microtubule connections during cell division. In striking contrast, burst size-dominated genes exhibited a distinct functional profile. Their shared GO terms across alleles primarily involve signaling transduction, gene regulation, energy/material metabolism, and cell structure/organelle organization. These findings reveal a clear functional partitioning: frequency-dominated regulation is almost exclusively dedicated to cell division processes, while burst size-dominated regulation governs signaling, metabolism, and structural organization. This specialization suggests that transcriptional control mechanisms have evolved to precisely coordinate gene expression for distinct biological tasks.

TFs are hypothesized to regulate burst kinetics [2, 5, 17]. For the C57 and CAST alleles, we performed TF enrichment analyses on gene populations with *b*_*f*_ -dominated and *b*_*s*_-dominated variations. Enriched TFs were categorized into three groups: the majority are general TFs that modulate both *b*_*f*_ and *b*_*s*_, while the less prevalent remaining two categories consist of specific TFs regulating only *b*_*f*_ or *b*_*s*_. Although specific TFs account for only a small proportion of total enriched TFs, they represent nearly 70% of the top ten TFs ranked by enrichment fold for either *b*_*f*_ or *b*_*s*_, underscoring their significant regulatory role. Notably, top-ranked TFs align with functions identified via GO term enrichment. For example, in *b*_*f*_ -dominated variant genes, *Foxo4, E2f2* in C57 [37, 38], *Rb1, Rbl2* in CAST [39, 40], and shared *Ing1* in both alleles [41], directly regulate cell division processes. Meanwhile, enriched *Smarca2* in CAST may provide a novel mechanistic explanation for its role in regulating “mitotic DNA replication initiation” gene accessibility via chromatin remodeling pathway [42, 43]. In CAST allele’s *b*_*s*_-dominated variant genes, top enriched TFs form a sophisticated network fine-tuning signaling for gene regulation: *Brd3* recognizes activation marks (e.g., histone acetylation) to promote transcription [44]; *L3mbtl2* and *Smchd1* establish inhibitory chromatin environments [45, 46]; *Hexim1* mediates rapid transcriptional pausing through RNA polymerase regulation [47]; and *Hnrnpk* determines mRNA fate via processing and stability modulation [48]. Furthermore, both alleles share *b*_*f*_ -specific TFs *Thrb*, and *b*_*s*_-specific TFs *Zfp322a, Smchd1, Hnrnpk, L3mbtl2*, findings that complement current knowledge of TFs regulating burst frequency and size.

### Transcriptomic regulation of burst frequency and size in human hearts with or without hypertrophic cardiomyopathy

Hypertrophic cardiomyopathy (HCM) is a prevalent inherited cardiovascular disorder, often progressing to heart failure or even sudden death [49, 50]. As the most common cardiac genetic disease, it is characterized by abnormal cardiomyocyte hypertrophy and excessive cardiac fibrosis. While existing single-cell transcriptome studies on human cardiomyopathies have focused primarily on pathological and clinical phenotypes, research into the regulatory mechanisms underlying transcriptional bursting remains notably limited. Single-nucleus RNA sequencing (snRNA-seq) has proven effective in dissecting cellular heterogeneity in the adult human heart under physiological conditions [49, 50]. To address this knowledge gap, we performed snRNA-seq on cardiac tissues from Chinese HCM patients and healthy donors. Specifically, left ventricular tissues were collected from 2 HCM patients who underwent cardiac transplantation, while healthy myocardial tissues were obtained from 2 donor hearts that could not be transplanted due to time constraints (Methods and Materials). Both sets of samples were processed for snRNA-seq (Methods and Materials).

All samples were sequenced individually, and after rigorous quality control, 28,321 nuclei were retained, including 11,830 from HCM patients and 16,491 from healthy controls (Supplementary Materials, Fig. S2). Based on the expression of established lineage markers [51], we employed the Seurat algorithm to identify and cluster cells from both HCM and healthy samples [52], as illustrated in Fig. 4a and Supplementary Materials Fig. S3. Twelve distinct cell types were identified, among which vascular endothelial cells, pericytes, fibroblasts, and cardiomyocytes were the four most abundant-consistent with previous findings [50]. Each of these four cell types contained over 500 nuclei in both the HCM and healthy groups (Fig. 4b), a sample size we validated to ensure the reliable application of the effective telegraph model to their snRNA-seq data (Fig. 2f-g). Conclusions regarding the effective *b*_*f*_ -dominated and *b*_*s*_-dominated variants thus reflect their respective actual regulatory modes, and our discussion focuses on burst regulation within these four cell types (Tables S3-S6).

**Figure 4.**
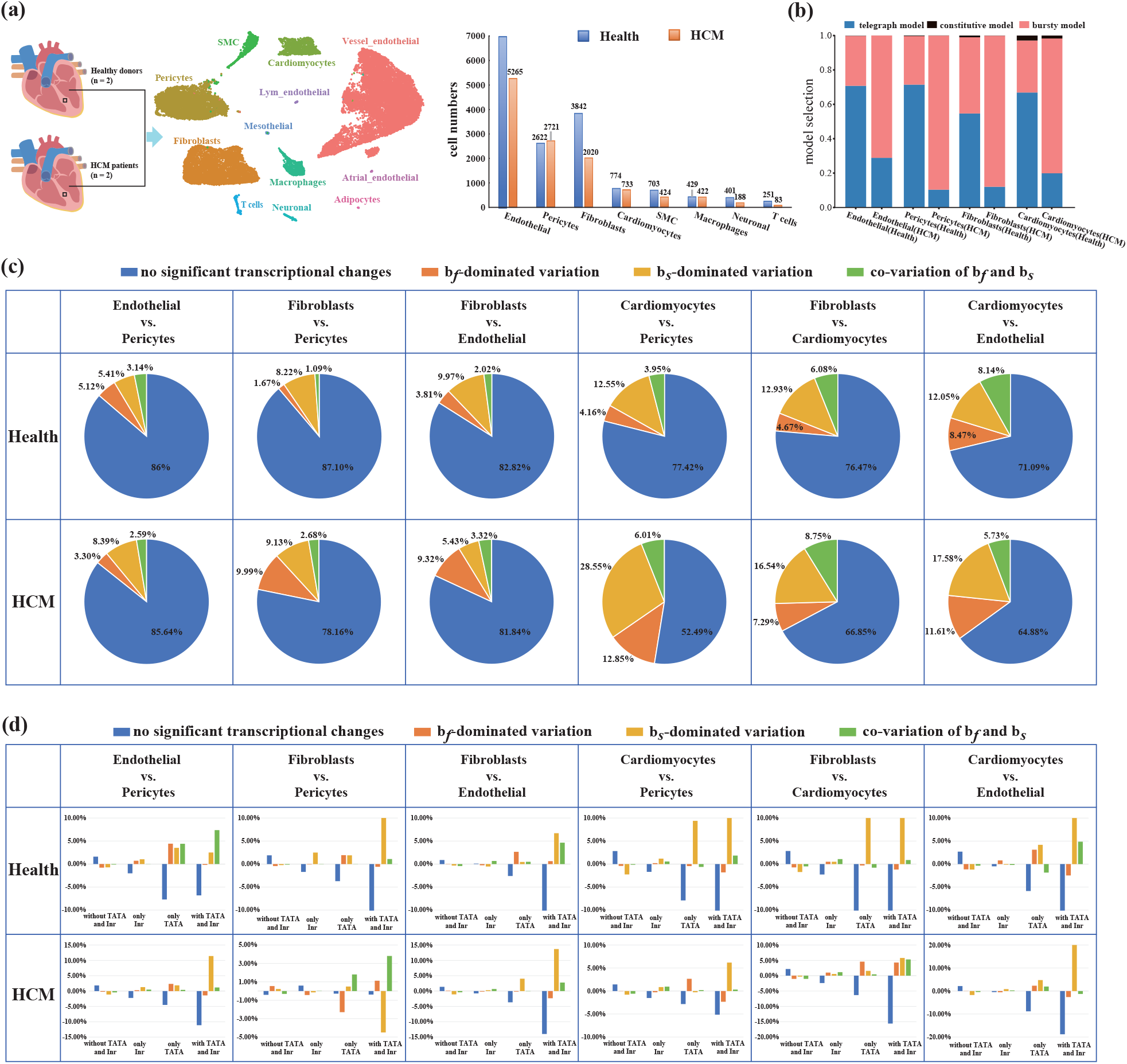
Transcriptional burst regulation in healthy vs. HCM human heart tissues. **(a)** Cell type identification via snRNA-seq: Left ventricular tissues (2 HCM patients and 2 healthy donors) yielded 28,321 qualified nuclei (HCM: 11,830; healthy: 16,491). Seurat clustering identified 12 cell types, with vascular endothelial cells, pericytes, fibroblasts and cardiomyocytes being the most abundant. **(b)** Model selection for gene product distributions: Constitutive models were rarely selected. Across the four key cell types, healthy samples had the highest proportion of genes described by the telegraph model, while HCM samples favored the bursty model. **(c)** *b*_*f*_ and *b*_*s*_ variation patterns across cell type pairs: Healthy samples exhibited the largest “no significant change” proportion; among significant changes, *b*_*s*_-dominated variation was most common. HCM samples showed reduced “no significant change” proportions and increased *b*_*f*_ - and *b*_*s*_-dominated variations. **(d)** Impact of promoter architectures (only TATA, only initiator, both, neither) across cell type pairs: Initiators had no effect; TATA elements independently reduced “no significant change” proportions. TATA-initiator synergy (observed in 9/12 cell type pairs) enhanced variations, primarily driving *b*_*s*_-dominated regulation.

We first examined differences in transcriptional bursting between HCM and healthy samples. The telegraph model has two special cases: the constitutive model, where an extremely high activation rate *b*_*f*_ = *k*_on_ keeps genes perpetually active, resulting in Poisson-distributed gene products [4, 17]; and the bursty model, characterized by extremely high inactivation rate *k*_off_ and synthesis rate *k*_b_, with a constant burst size *b*_*s*_ = *k*_b_*/k*_off_ within a normal range. In the bursty model, the gene remains in an inactive state for most of the time, but once activated, it produces a large number of products, generating a negative binomial distribution [53]. We applied the corrected Akaike information criterion (AICc) [21, 22] to the snRNA-seq data of the four cell types from both HCM and healthy samples (Methods and Materials). As shown in Fig. 4b, the constitutive model was rarely selected. Notably, across all four cell types, HCM samples had the highest proportion of genes best described by the bursty model, whereas healthy samples had the highest proportion best fit by the telegraph model. These results demonstrate that transcriptional bursting is significantly more prevalent in diseased cardiomyocytes compared to healthy ones, underscoring the need for further analysis of its regulation.

Next, we examined changes in effective *b*_*f*_ and *b*_*s*_ across different cell type contexts (Fig. 4c). For both HCM and healthy samples, we analyzed the 6 pairwise combinations of the four cell types. Herein, we focused genes best characterized by the telegraph or bursty models, while excluding those with a fold change in *b*_*f*_ (and *b*_*s*_) ratios exceeding 1000 (Table S7). In healthy samples, variations in *b*_*f*_ and *b*_*s*_ between these 6 cell type pairs fell into four patterns: *b*_*f*_ -dominated variation, *b*_*s*_-dominated variation, co-variation, and no significant change-consistent with our analysis of mouse fibroblasts and ES cells (Fig. 3a). Notably, the “no significant change” pattern accounted for the largest proportion of genes in healthy samples. Among genes with significant changes, however, *b*_*s*_-dominated variation was most prevalent, in contrast to the mouse model where *b*_*f*_ -dominated variation was the primary type. In HCM samples, we observed a similar proportional distribution of these variation patterns but with a key difference: for each cell type pair, the proportion of genes with no significant changes in *b*_*f*_ and *b*_*s*_ was significantly lower than in healthy samples. This reduction was accompanied by a marked increase in genes with regulatory changes in *b*_*f*_ and *b*_*s*_ in HCM samples, with particularly notable rises in the proportions of *b*_*f*_ -dominated and *b*_*s*_-dominated variations. These findings indicate a shift toward more dynamic and widespread transcriptional burst regulation in HCM, which may play a role in driving its pathological phenotypes.

We further explored how promoter TATA and initiator elements influence variations in effective *b*_*f*_ and *b*_*s*_, analyzing 12 cell type pairs from HCM and healthy samples (Fig. 4d). Following the approach for mouse fibroblasts and ES cells (Fig. 3b), we categorized genes into four promoter groups (TATA-only, initiator-only, both, or neither) and subdivided each into four *b*_*f*_ and *b*_*s*_ variation categories (no change, *b*_*f*_ -dominated, *b*_*s*_-dominated, co-variation). Regulatory impact was assessed by comparing group proportions to baseline (unstratified by promoters in Fig. 4c). Consistent with findings in the mouse model (Fig. 3b), promoters lacking both elements or with only initiators had no significant effect on genes with unchanged *b*_*f*_ and *b*_*s*_. In contrast, TATA-only promoters significantly reduced this “no change” proportion, confirming their independent role in promoting *b*_*f*_ and *b*_*s*_ variations-mirroring the mouse model. Notably, in 9 of 12 pairs, promoters with both elements exerted marked synergy, further reducing the “no change” proportion, enhancing *b*_*f*_ and *b*_*s*_ variations, and primarily driving *b*_*s*_-dominated changes. These cross-species results reveal a unified model: initiators have no significant impact, TATA elements independently promote *b*_*f*_ and *b*_*s*_ variations, and their synergy specifically drives *b*_*s*_-dominated regulation in both HCM and healthy samples.

### Lineage-specific regulatory changes: burst frequency- and size-dominated drivers of cardiomyocyte hypertrophy

In this section, we examine burst regulatory alterations between HCM and healthy samples across four distinct cell types, focusing on four primary patterns: *b*_*f*_ -dominated (burst frequency-dominated), *b*_*s*_-dominated (burst size-dominated), co-variation, and no significant change. As shown in Fig. 5a, among endothelial cells, only 3% of genes exhibit significant changes in *b*_*f*_ or *b*_*s*_, indicating that gene transcriptional regulation in these cells remains relatively stable during the progression of hypertrophic cardiomyopathy. In contrast, pericytes and fibroblasts separately show an 8% and 11% proportions of genes with significant changes in either *b*_*f*_ or *b*_*s*_, with a predominance of *b*_*s*_-dominated and *b*_*f*_ -dominated alterations. This observation aligns with prior literature highlighting the involvement of pericytes and fibroblasts in extracellular matrix remodeling and paracrine signaling during HCM pathogenesis [50]. Notably, cardiomyocytes display the largest proportion of significant *b*_*f*_ and *b*_*s*_ changes, approaching 30%, which further confirms their central role in driving the hypertrophic response through dysregulated transcriptional dynamics [49, 50]. In summary, these lineage-specific regulatory changes underscore cell-type-specific contributions to HCM pathogenesis, with cardiomyocytes emerging as primary drivers of transcriptional burst dysregulation, endothelial cells maintaining relative stability, and fibroblasts showing intermediate levels of transcriptional remodeling.

**Figure 5.**
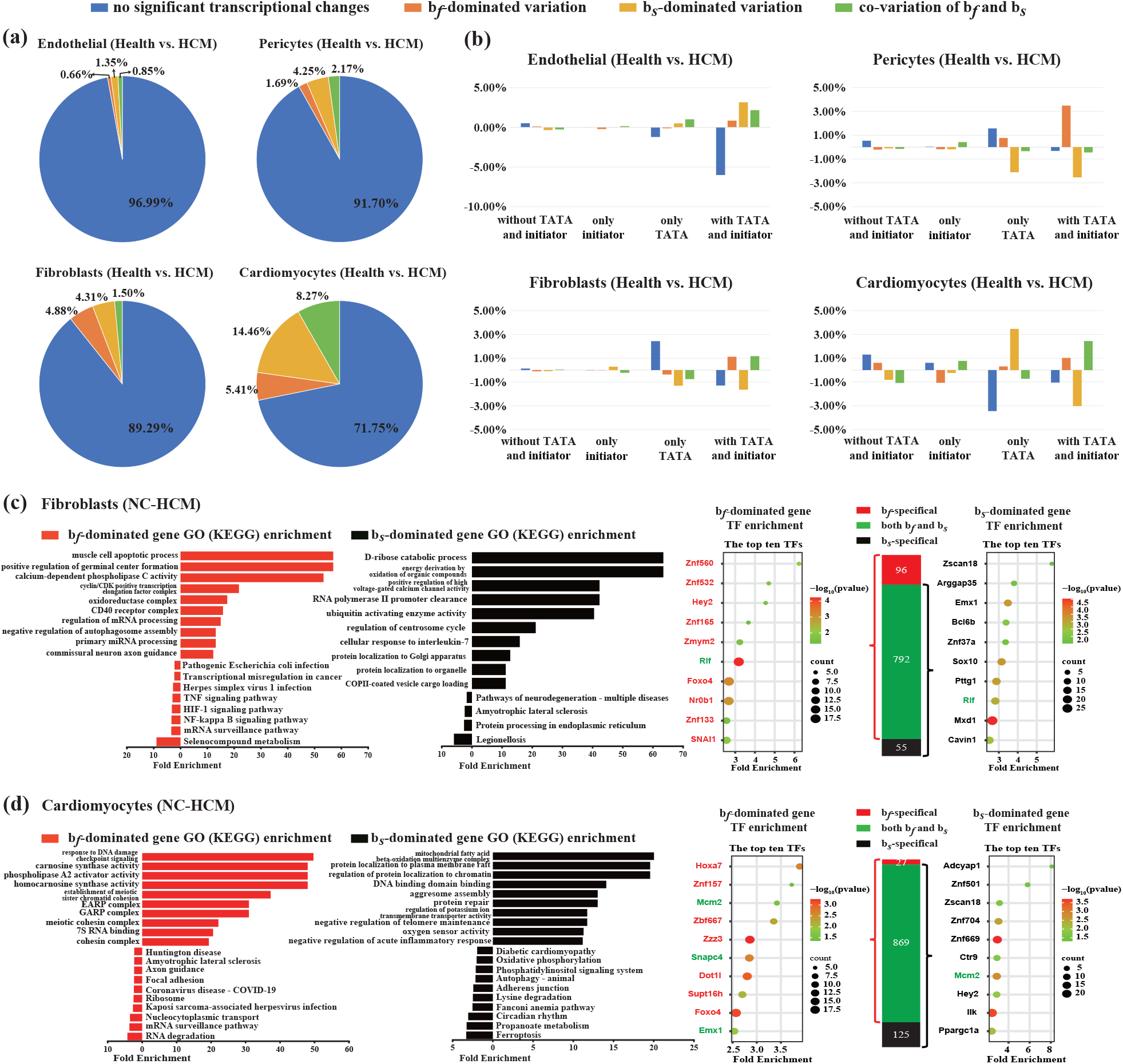
Cell-type-specific mechanistic analysis of transcriptional burst regulation in HCM. **(a)** Burst parameter variation patterns in four heart cell types (HCM vs. healthy): Endothelial cells had minimal significant *b*_*f*_ and *b*_*s*_ changes (3% of genes); cardiomyocytes showed the most pronounced dysregulation (30% of genes with significant *b*_*f*_ and *b*_*s*_ changes); Pericytes and fibroblasts displayed intermediate burst parameter changes. **(b)** The TATA-Initiator synergy enhancing *b*_*f*_ and *b*_*s*_ variations was retained only in stable endothelial cells, while absent in pericytes, fibroblasts, and cardiomyocytes in the context of HCM compared to healthy tissues. **(c)-(d)** GO, KEGG pathway, and TF enrichment analyses were conducted for *b*_*f*_ - and *b*_*s*_-dominated genes in fibroblasts (c) and cardiomyocytes (d) to identify their biological functions and regulatory mechanisms in HCM vs. healthy comparisons.

Previously, across different cell types, we found initiators exert no significant impact on *b*_*f*_ and *b*_*s*_, while TATA elements independently drive *b*_*f*_ and *b*_*s*_ variations; their synergy specifically orchestrates *b*_*f*_ - or *b*_*s*_-dominated regulation in mouse cells (Fig. 3b) and human heart tissue (Fig. 4d). Here, we extend this analysis to four major cell types-endothelial cells, pericytes, fibroblasts, and cardiomyocytes-to explore whether this regulatory pattern persists under healthy versus pathological (HCM) conditions. Across all four cell types, initiators remain without significant impact, and TATA elements lack consistent independent effects on *b*_*f*_ and *b*_*s*_ variations (Fig. 5b). The key divergence lies in the synergistic effect of TATA and initiator elements: it is preserved only in endothelial cells, where pathological environments do not significantly alter *b*_*f*_ and *b*_*s*_ regulation (Fig.,5a). In pericytes, fibroblasts, and cardiomyocytes-where *b*_*f*_ and *b*_*s*_ undergo robust pathological alterations-this synergism is lost. This loss is likely driven by pathological environmental cues (e.g., heightened cellular stress signaling and remodeled epigenetic marks), which override the intrinsic regulatory contributions of TATA and initiator elements.

We further focused on exploring the pathological associations of burst frequency-dominated (*b*_*f*_ -dominated) and burst size-dominated (*b*_*s*_-dominated) regulatory genes in cardiomyocytes and fibroblasts, guided by their central relevance to HCM. As the most prevalent cardiac genetic disorder, HCM is defined by two hallmark pathologies: cardiomyocyte hypertrophy and excessive cardiac fibrosis-processes driven primarily by cardiomyocytes and fibroblasts, respectively [49, 50]. Notably, these two cell types also exhibit the most pronounced shifts in *b*_*f*_ and *b*_*s*_ regulation when comparing healthy and HCM samples (Fig. 5a), making them ideal targets for investigating how transcriptional burst dynamics contribute to disease progression.

In fibroblasts from healthy versus HCM samples, functional enrichment analyses revealed distinct roles for *b*_*f*_ - and *b*_*s*_-dominated regulatory genes (Fig. 5c). GO enrichment analysis of *b*_*f*_ -dominated differential genes highlights muscle cell apoptotic process as significantly enriched pathways. Notably, TF *Foxo4* is identified as one of the top regulator in these genes. KEGG analysis further underscores the significance of NF-*κ*B, TNF, and HIF-1 signaling pathways. For *b*_*s*_-dominated differential genes, GO enrichment highlights pathways linked to energy metabolism including D-ribose catabolic process and energy derivation by oxidation of organic compounds (Fig. 5c). Notably, protein processing in endoplasmic reticulum pathway is significantly enriched in bs dominated genes.

In cardiomyocytes, functional enrichment analyses also revealed distinct roles for *b*_*f*_ - and *b*_*s*_-dominated regulatory genes (Fig. 5d). *b*_*f*_ -dominated genes were significantly enriched in GO terms associated with response to DNA damage checkpoint signaling–a pathway critical for maintaining genomic stability. Additionally, KEGG analysis further underscores that *b*_*f*_ -dominated genes have showed enrichment in ribosome pathways. *b*_*s*_-dominated genes in cardiomyocytes are associated with a broader network of dysregulated pathways, including metabolic, post-transcriptional, and cell death processes (Fig. 5d). GO analysis reveals strong enrichment in mitochondrial fatty acid *β*-oxidation multienzyme complex and protein localization to plasma membrane raft. Central to this axis is the top enriched TF *Ppargc1a* (PGC-1). KEGG pathway analysis further highlighted the enrichment of propanoate metabolism and oxidative phosphorylation pathways, demonstrating disturbances in energy metabolism within the myocardium. Notably, these findings are also linked to ferroptosis.

## Conclusions and discussions

Gene transcription is inherently a stochastic bursting process, with burst frequency and burst size serving as core parameters that govern the dynamics of gene expression [2, 6]. Current discussions on stochastic gene expression models have two key limitations. First, while the classical telegraph model is widely used to fit single-cell transcriptional data for its computational stability and focus on two-state transitions (between active and inactive states) [4, 6, 8], it overlooks the intricate mechanisms governing genome-wide genes, such as three-state models with multi-step activation [26, 27], cross-talk pathway models with competitive signaling [28, 29], and feedback models with autoregulatory circuits [31, 32]. This raises a critical question: does telegraph model-inferred burst regulation truly reflect actual regulatory dynamics? Second, despite growing focus on more complex systems, our work confirms that classical inference methods (e.g., AICc) and maximum likelihood estimation are unstable for models with ≥ 3 parameters [22]. Existing theoretical results rely heavily on large samples (*N* = 10^4^) [36], time-dependent data for kinetic informations [23], and validation across diverse data types [54]. Computationally, these approaches often require error-prone approximations [55] or simplifications to reduce model complexity [25]. While feasible for specific genes, these strategies struggle to generalize to steady-state scRNA-seq and smFISH distribution data, limiting genome-wide utility.

In this study, we employ the simple telegraph model to analyze transcriptional burst regulation across genome-wide scRNA-seq and smFISH datasets, addressing critical gaps in understanding and interpreting bursting dynamics in complex biological systems. While the telegraph model exhibits quantitative biases in estimating its effective burst frequency *b*_*f*_ and size *b*_*s*_ compared to actual values for genes under complex regulatory frameworks (e.g., three-state, cross-talk pathway, or feedback models), it reliably captures qualitative changes in burst dynamics. Specifically, with sample sizes *N* ≥ 500 and a 3-fold change in *b*_*f*_ or *b*_*s*_, variations inferred as *b*_*f*_ -dominated and *b*_*s*_-dominated from the telegraph model serve as robust proxies for true frequency-driven and size-driven regulatory changes, respectively. This robustness enables its application to genome-wide datasets where fitting complex models for each gene is impractical. Transcriptomic analyses using the telegraph model reveal three general rules for the functional specialization of burst frequency-dominated and size-dominated regulation across diverse contexts, including mouse CAST and C57 alleles (encompassing fibroblasts and ES cells) and human heart tissues under both healthy and HCM conditions (encompassing vascular endothelial cells, pericytes, fibroblasts and cardiomyocytes).

First, among genes subject to burst regulation, the majority exhibit either burst frequency-dominated or burst size-dominated patterns. This observation holds consistently across various contexts: comparisons between mouse fibroblasts and ES cells (both CAST and C57 alleles); between different human heart cell types (under healthy and HCM conditions); and within the same cell types when comparing healthy and pathological states. These findings aligns with existing literature, which documents similar regulatory patterns across genes under diverse promoters, chromatin structures, and external stimuli. Examples include the varied burst frequency of mouse genes during the cell cycle [14], elevated burst size in over 20 *E. coli* promoters [15], and the serum-induced regulation of mammalian *ctgf* gene [16].

Notably, a small proportion of burst regulation exhibits *b*_*f*_ and *b*_*s*_ co-variation. This phenomenon may reflect genuine coordinated regulation of burst frequency and size or, alternatively, regulation of a single parameter within a more complex system. Distinguishing between these underlying mechanisms typically requires additional temporal information [21, 23]. However, certain cases of co-variation can be readily explained. For instance, the regulatory patterns of the HIV LTR promoter across human chromatin environments and the regulation of gap genes along the anterior-posterior axis in Drosophila embryos align with the telegraph model: targeting *b*_*f*_ at low expression levels and *b*_*f*_ at high expression levels [3, 7]. This corresponds to the observed regulatory pattern where burst frequency dominates at low expression and burst size dominates at high expression levels [5].

Second, in regulating burst frequency and size, promoter initiator elements are nearly ineffective, while TATA elements drive such regulation; together, they exhibit a synergistic effect that primarily promotes burst size-dominated regulation. This pattern, however, is confined to environmental variations across different cell types in mouse and human cardiac tissues. Within the same cell type, this regulation-particularly the synergistic effect-vanishes when transitioning from healthy to pathological HCM conditions. Our results validate prior findings that TATA boxes promote burst size [2, 6, 12, 13] and initiators enhance this effect [2, 6] in human and mouse cells. Notably, while existing studies focus on TATA box and initiator regulation of bursting across different genes within a single cell type, we explore their modulation of the same gene under distinct cellular environments. A striking observation is the complete loss of TATA-initiator synergy in healthy versus HCM samples, likely due to pathological cues inducing alternative regulatory modes that overshadow promoter regulation. Indeed, diverse TATA-independent modes exist, including *Med26*’s profound impact on gene regulatory networks downstream of chromatin architecture [13], Akt/MAPK pathway modulation via transcriptional elongation efficiency [12], and enhancers’ critical roles in shaping transcriptional mechanisms [2].

Third, burst frequency- and size-dominated regulation are linked to distinct gene functions; top TFs governing these modes are mostly specific, regulating either frequency- or size-dominated variation, but not both. In a comparison of mouse fibroblasts versus ES cells, frequency-dominated differentially expressed genes (DEGs) were significantly enriched in core cell division events. This highlights that frequency variations primarily impact cell cycle precision and genome inheritance fidelity, aligning with prior findings in mouse ES cells where *Oct4* and *Nanog* regulate burst frequency post-gene replication [14]. Notably, frequency-enriched TFs differ between C57 and CAST alleles but modulate similar biological functions via distinct pathways. For example, in C57, *E2f2* controls the G1/S transition via the *Rb/E2f* pathway, directly activating DNA replication genes [37]; in CAST, *Rbl2*, a key regulator in the same pathway, maintains cell cycle checkpoint function by inhibiting *E2f* activity [40]. These regulatory differences likely represent adaptive cell cycle network variations across genetic backgrounds to support the fibroblast-to-pluripotency transition. In contrast to the highly focused regulation of burst frequency, burst size-dominated DEGs in mouse fibroblasts versus ES cells exhibit diverse functions across alleles. GO enrichment analysis reveals functions such as signaling, gene regulation, metabolism and cell structure. This indicates that burst size variations more broadly modulate basal cellular physiology and functional traits.

In fibroblasts from healthy versus HCM samples, burst frequency-dominated DEGs are enriched in pathways regulating cell apoptosis, with *Foxo4* identified as a key TF known to regulate these processes [56]. *Foxo4* is known to control both autophagy and apoptosis, suggesting its potential role in driving these dysregulated processes in HCM [56]. Apoptosis, a tightly regulated process of programmed cell death, contributes to myocardial cell depletion in various cardiovascular diseases, such as myocardial infarction, ischemic heart disease, and heart failure [57]. Dysregulated apoptosis is associated with adverse cardiac outcomes [57, 58]. KEGG analysis further reveals frequency-related enrichment in NF-*κ*B, TNF, and HIF-1 signaling pathways, which are key cascades activated in oxidative stress, inflammation, and hypoxia-hallmarks of cardiovascular diseases [59]. In contrast, burst size-dominated DEGs are primarily enriched in energy metabolism pathways, indicating oxidative stress and energy metabolic dysfunction in HCM fibroblasts-phenotypes strongly associated with heart failure development and consistent with extensive studies linking mitochondrial dysfunction to heart failure [60]. Additionally, these DEGs cluster in endoplasmic reticulum protein processing pathways, underscoring the clinical relevance of protein homeostasis disruption, a known driver of multiple forms of heart disease [61]. Collectively, these findings indicate that aberrant transcriptional regulation and disrupted protein homeostasis jointly promote fibroblast activation and cardiac remodeling in cardiomyopathy, offering potential multi-target therapeutic strategies to alleviate cardiac fibrosis progression.

In cardiomyocytes from healthy versus HCM samples, burst frequency-dominated DEGs are enriched in DNA damage checkpoint signaling, a critical pathway for maintaining genomic stability [62]. Dysregulation in this pathway may be driven by key TFs such as *Foxo4* (a stress-responsive regulator interacting with DNA repair genes) [63] and *Mcm2* (a core factor in preserving genomic replication stability) [64], potentially leading to accumulated DNA damage, reduced genomic stability, and subsequent cardiomyocyte death. Additionally, ribosomal pathway enrichment suggests altered ribosomal profiles by differential expression of ribosomal proteins (e.g., *Rpl30, Rps27, Rpl21*). This could result in insufficient production of essential cardiac proteins or accumulation of misfolded proteins, further impairing myocardial repair [65], and may reflect a potential mechanism for switching between basal cardiac function and stress-induced recovery modes [66]. In contrast, burst size-dominated DEGs are enriched in mitochondrial fatty acid *β*-oxidation pathways and protein localization to plasma membrane raft, both of which are critical to cardiac energy metabolism and membrane homeostasis. Mitochondrial dysfunction, a well-established pathogenic driver in HCM and heart failure [67], is likely exacerbated by this impaired fatty acid oxidation, which directly compromises cardiac contractility [68]. The top TF *Ppargc1a*, a master regulator of mitochondrial function [69], is likely mediating metabolic dysregulation in this context. KEGG analysis further reveals perturbations in ferroptosis (a process linked to cardiovascular disease [70]) and propionate metabolism, reflecting energy dysfunction. Notably, these burst frequency-dominated and size-dominated regulatory axes are interdependent. Genomic instability resulting from abnormal burst frequency regulation directly impairs mitochondrial function, while metabolic dysfunction arising from burst size dysregulation exacerbates oxidative stress, further damaging DNA and proteins. This creates a vicious cycle that accelerates myocardial degeneration.

In summary, this study establishes the telegraph model as a practical framework for decoding genome-wide transcriptional burst regulation. Linking burst parameters to biological function and disease, it elucidates how gene expression stochasticity sustains cellular homeostasis and is disrupted in pathological states, while paving the way for targeted burst manipulation in regenerative medicine and therapy. A key limitation of this study is the telegraph model’s failure to fully capture quantitative features of complex regulatory mechanisms. Future work should incorporate additional regulatory mechanisms into computationally scalable frameworks, potentially integrating machine learning and stochastic modeling to improve parameter inference efficiency [24, 25]. Furthermore, while the current analyses focus on steady-state data, coupling transcriptional burst dynamics with time-resolved single-cell measurements (e.g., nascent RNA live imaging) could provide deeper insights into how burst parameters respond to acute stimuli [23, 54]. Another promising avenue is to explore differential expression analysis in scRNA-seq data based on the telegraph model’s characterization of burst frequency and size [8, 25]. This mechanistic framework could help link a gene’s functional role to a cell’s regulatory strategies, moving beyond observational changes in mean expression. Such analyses may reveal how transcriptional burst dynamics evolve to underpin species-specific traits or disease susceptibilities.

## Methods and Materials

### Computation of burst frequency and burst size for each model

For convenience, we set the degradation rate *k*_d_ = 1 for all models. This choice is justified by the fact that parameters-with the exception of *q*_1_ and *q*_2_ in the cross-talk pathway model-can always be non-dimensionalized using *k*_d_ for computing the steady-state gene product distributions [21]. Thus, all parameters presented in this study (excluding *q*_1_ and *q*_2_ should be interpreted as their actual values normalized by *k*_d_. This assumption also implicitly assumes a consistent *k*_d_ across different cell types [6, 8, 12, 13]: when calculating the ratios of two *b*_*f*_ (or *b*_*s*_) values, *k*_d_ is automatically cancelled out.

Across all tested models, *k*_b_ denotes the synthesis rate of the gene product. Let *T*_on_ and *T*_off_ stand for the mean active and inactive durations of the gene, respectively. Burst frequency is thus calculated as 1*/T*_off_, while burst size is determined as *k*_b_*T*_on_. For the telegraph model, *T*_on_ = *k*_off_ and *T*_off_ = *k*_on_ [53]. For the three-state model, gene inactivation involves a single exponential step, whereas gene activation comprises two exponential steps-allowing straightforward calculation of *T*_on_ and *T*_off_ as follows [26]:

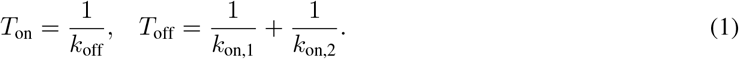

For the cross-talk pathway model, gene inactivation occurs via one pathway, while gene activation proceeds through two pathways; this enables simple computation of the mean active and inactive durations [28]:

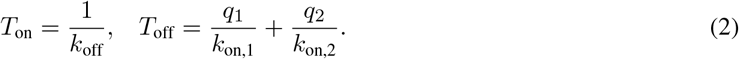

We next calculate *T*_on_ and *T*_off_ for the two feedback models. Reference [32] demonstrates that regardless of whether autoregulation is present, the mean ⟨*m*⟩ of gene product counts can consistently be expressed in terms of the mean active and inactive periods as:

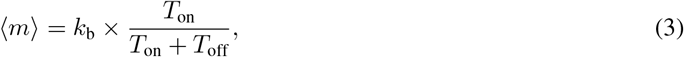

where the second term on the right-hand side represents the probability of the gene being in an active state. The exact formulas for ⟨*m*⟩ involve complex hypergeometric functions (Supplementary Material). To mitigate computational errors associated with calculating these hypergeometric functions, we determine ⟨*m*⟩ by first applying the finite-state projection (FSP) algorithm to compute the gene product distribution, then calculating the mean of this distribution (see online code). For the positive feedback model, the mean active duration is given by *T*_on_ = 1*/k*_off_. Substituting this expression into Eq. (3) yields an explicit formula for the mean inactive duration:

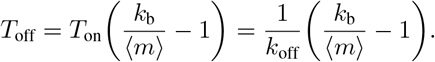

Similarly, for the negative feedback model, the mean inactive duration is given by *T*_off_ = 1*/k*_on_. Substituting this into Eq. (3) produces an explicit expression for the mean active duration:

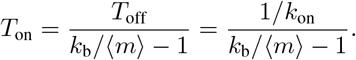

### Parameter selection for each model

In Figs. 1-2, we generated synthetic gene product number data under different parameter sets for the simple telegraph model and four complex models. For the telegraph model, we randomly selected parameters as follows: *k*_on_ = 1*/T*_off_ ∈ [0.1, 5], *k*_off_ = 1*/T*_on_ ∈ [0.1, 5] and *k*_b_ ∈ [5, 50]-values that span a broad range of parameter spaces [1, 2, 6, 7, 15, 17, 53]. For the three-state and cross-talk pathway models, *k*_b_ and *k*_off_ were randomly selected using the same ranges as in the telegraph model. For other parameters, we first randomly set 1*/T*_off_ ∈ [0.1, 5]. For the three-state model, *k*_on,1_ was randomly chosen within [1*/T*_off_, 5]; according to Eq. (1), we can then write the expression for *k*_on,2_ = 1*/*(*T*_off_ − 1*/k*_on,1_); For the cross-talk pathway model, we first randomly selected *q*_2_ (or *q*_1_) ∈ (0, 1) and *k*_on,1_ ∈ [1*/T*_off_, 5]; according to Eq. (2), we can then write the expression for *k*_on,2_ = *q*_2_*/*(*T*_off_ − *q*_1_*/k*_on,1_). For the positive feedback model, *k*_b_ and *k*_off_ follow the same random selection ranges as the telegraph model. *k*_on_ is randomly chosen from [0.1, 3] and positive feedback rate *µ* from [0.05, 2], which ensures its *T*_off_ roughly lies within [0.1, 5]. For the negative feedback model, *k*_b_ and *k*_on_ adopt the same random selection ranges as the telegraph model. *k*_off_ is randomly selected from [0.1, 3] and negative feedback rate *ν* from [0.05, 1], forcing its *T*_on_ to be roughly within [0.1, 5].

### Parameter inference for the telegraph model (maximum-likelihood estimation method)

We use the “fminsearch” function in MATLAB to minimize the negative log-likelihood function, defined as:

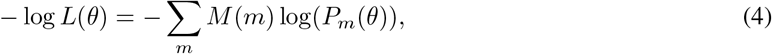

where *θ* denotes the parameter set of the model, *M* (*m*) is the number of cells containing *m* copies of the gene product, and *P*_*m*_(*θ*) represents the gene product distribution parameterized by *θ*. Through this minimization, we obtain estimates of the three telegraph model parameters *k*_on_, *k*_off_ and *k*_b_, with *k*_d_ = 1 as established earlier.

A critical step in this optimization is the selection of initial values 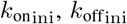, and 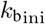 for *k*_on_, *k*_off_ and *k*_b_, respectively. A modified moment estimation method has been proposed [21]: 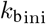 is typically set to the maximum number of gene product molecules observed in the cell population, and 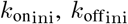 are reversely derived directly from the analytical formulas for the mean and Fano factor (Variance over mean) of the gene product distribution in the telegraph model. However, for real smFISH or scRNA-seq datasets, this approach often yields excessively large 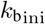 values for certain genes, drastically slowing optimization. Furthermore, the Fano factor of the distribution data may be < 1-contradicting the inherent property of the telegraph model, whose Fano factor is always > 1. Thus, we did not use the moment-based method to determine initial parameters in this study. Instead, we performed 20 independent minimization runs for each optimization, using randomly generated initial parameters 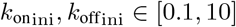 and 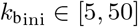 [6, 53].

### Model Selection: constitutive, bursty, and telegraph models

The constitutive model is a special case of the telegraph model, defined by an extremely high activation rate *k*_on_ ≫ 1. This leads to Poisson-distributed gene products, which are characterized by a single parameter *k*_b_ [4]. Another special case of the telegraph model is the bursty model, defined by extremely high synthesis and inactivation rates *k*_b_, *k*_off_ ≫ 1. This results in a negative binomial distribution, described by two parameters *k*_on_ and *k*_b_*/k*_off_ [53]. For experimental steady-state mRNA number distributions, we fit the data to the constitutive, bursty, and telegraph models separately by minimizing the negative log-likelihood function log *L*(*θ*) defined in (4). To identify the best model for explaining the distribution data, a common approach is to select the one with the lower corrected Akaike information criterion (AICc)-a metric that incorporates penalties for the number of parameters and sample size [21, 22]:

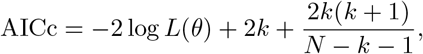

where *k* is the number of model parameters (*k* = 1 for the constitutive model, *k* = 2 for the bursty model, *k* = 3 for the telegraph model) and *N* is the cell sample size.

### Human heart tissue collection

Left ventricular tissues were collected from 2 patients diagnosed with hypertrophic cardiomyopathy (HCM group) who underwent cardiac transplantation at Guangdong Provincial People’s Hospital, Guangzhou, China.

Healthy left ventricular myocardial tissues (healthy group, *n* = 2) were obtained from donor hearts that could not be transplanted due to timing constraints. All excised heart tissues were immediately flash-frozen and stored in liquid nitrogen until further analysis. This study was carried out in compliance with the Declaration of Helsinki and approved by the Ethics Committee of Guangdong Provincial People’s Hospital (GDREC2016255H). Written informed consent was obtained from all participants.

### snRNA-sequencing

The cell suspension was resuspended in Labeling Buffer with PBS containing 0.04% BSA. The prepared samples were measured by Fluorescence Cell Analyzer (countstar) for cell count and viability. After using MACS Dead Cell Removal Kit (Miltenyi, 130-090-101) to remove dead cells in samples with low activity according to the instructions, using a Single Cell 3’ Library and Gel Bead kit (10X Genomics, cat 1000092) and Chromium Single Cell B Chip kit (10X Genomics, cat 1000074), the cell suspension was loaded onto a Chromium single cell controller (10X Genomics) to generate single-cell gel beads in the emulsion (GEMs). Captured cells were lysed and the released RNA was barcoded through reverse transcription in individual GEMs. Reverse transcription was performed on a C1000TM Touch Thermal Cycler (Bio Rad) at 53^*°*^C for 45min, followed by 85^*°*^C for 5 min and a hold at 4^*°*^C. Complementary DNA was generated and amplified, after which, library was constructed with Library Construction Kit(10x Genomics, cat 1000190).

The libraries were sequenced using an Illumina Hiseq sequencer with a paired-end 150-bp (PE150) reading strategy.

Single-cell RNAseq data were processed using the Cell Ranger (v.3.0.1) with GRCh38 human reference.

## Data accessibility

The transcriptomic data from both healthy human and HCM heart samples, generated in this study, have been deposited in the National Genomic Data Center (NGDC) database under the accession number HRA013899, and access to the data requires formal approval.

## Code accessibility

MATLAB code encompassing the Stochastic Simulation Algorithm (SSA), Finite-State Projection (FSP) algorithm, parameter inference (via maximum-likelihood estimation, MLE) and model selection (via corrected Akaike information criterion, AICc), and the Seurat algorithm used for identifying and clustering cells from human heart samples is available at https://github.com/Always-Stude/code.

The recognition information and coordinates of promoter motifs (TATA box, Initiator) were downloaded from the Eukaryotic Promoter Database (EPD). The database is accessible at: https://epd.expasy.org/epd/EPDnew_select.php.

Functional enrichment analysis (GO, KEGG, TFs) was performed using DAVID (The Database for Annotation, Visualization, and Integrated Discovery). The database is accessible at: https://davidbioinformatics.nih.gov/home.jsp.

## Author Contributions

Yueheng Wu, Guozhi Jiang, Jianshe Yu, and Feng Jiao conceived the original study. Liang Chen and Yu Liao performed theoretical derivations. Liang Chen, Yuwen Wu, and Kai Chen performed numerical simulations and data analysis. Liang Chen, Yuwen Wu, Sijia Fang, Guozhi Jiang, and Feng Jiao interpreted the theoretical results. Feng Jiao and Jianshe Yu provided supervision. Guozhi Jiang, Jianshe Yu, and Feng Jiao were responsible for project administration. Guozhi Jiang, Jianshe Yu, and Feng Jiao jointly wrote the manuscript with assistance from the co-authors.

## Acknowledgments

This work was supported by grants from the Natural Science Foundation of China (No. 12271118, 12331017, 82574188), Guangdong Basic and Applied Basic Research Foundation (No. 2024A1515013260), Guangzhou Municipal Science and Technology Plan Basic and Applied Basic Research (No. 2025A03J3089).

